# Tremors in Time: Mechanically Induced Motor Tremors Influence Time Perception

**DOI:** 10.1101/2023.08.01.551412

**Authors:** Keri Anne Gladhill, Rose De Kock, Weiwei Zhou, Wilsaan Mychal Joiner, Martin Wiener

## Abstract

Contemporary research has begun to show a strong relationship between movements and the perception of time. More specifically, concurrent movements serve to both bias and enhance time estimates. To explain these effects, we recently proposed a mechanism by which movements provide a secondary channel for estimating duration that is combined optimally with sensory estimates, in accordance with Bayesian cue combination. However, a critical test of this framework is that by introducing “noise” into movements, sensory estimates of time should similarly become noisier in a manner predicted by cue combination equations. To accomplish this, we had human participants move a robotic arm while estimating intervals of time in either auditory or visual modalities (n=24, ea.). Crucially, we introduced an artificial “tremor” in the arm while subjects were moving, that varied across three levels of amplitude (1-3 N) or frequency (4-12 Hz). The results of both experiments revealed that increasing the frequency of the tremor led to noisier estimates of duration, but in such a way that higher levels of noise saturated the impact, consistent with optimal integration. Further, the effect of noise varied with the base precision of the interval, such that a naturally less precise timing (i.e. visual) was more influenced by the tremor than a naturally more precise modality (i.e. auditory). To explain these findings, we fit the data with a recently developed drift-diffusion model of perceptual decision making, in which the momentary, within-trial variance was allowed to vary across conditions. Here, we found that the model could recapitulate the observed findings, further supporting the theory that movements influence perception directly. Overall, our findings support the proposed framework, and demonstrate the utility of inducing motor noise via artificial tremors, thus providing clinical utility in their connection to movement disorders characterized by tremors.

## Introduction

An abundance of previous research indicates that movement parameters differentially affect time estimates - in some cases movement biases the perception of time whereas in other cases it improves the precision of time estimates. It has recently been suggested that these contrasting effects can be explained via a Bayesian cue combination framework in which, along with sensory input such as auditory and visual information, movement itself also serves as an additional input for duration information (De Kock et al., 2021).

Evidence for this framework comes from work over the past decade demonstrating the impact of movements on time estimates. In particular, we have shown that, when subjects are allowed to freely move a robotic arm, as opposed to having the arm restrained in place, their estimation of concurrently presented sensory time estimates is more precise (Wiener et al., 2019). This effect occurs both within and between subjects, across different task designs, and does not depend on the type of movement strategy employed. Rather, the effect appears to depend on whether the subject is *moving or not*. In a further series of experiments, we additionally found that increasing the viscosity of movements, such that movement lengths were shortened, also shortened time estimates (De Kock et al., 2021). Further, this effect was tied to changes in the perception of duration, rather than to biases in decision making.

The cue combination framework proposes a manner in which noisy estimates are combined optimally by shifting the temporal estimates toward their more precise input (Ernst and Banks, 2002; Alais and Burr, 2019); therefore, since movements have been shown to be precise, with low variability and high temporal fidelity, the variance of movement time estimates will also be low (Brenner et al., 2012; Doumas et al., 2008). However, the brain also integrates statistics of body movements such as the speed, length, direction, and area of movement with other sensory input (Petzschner et al., 2015) leading to biases in the perception of time. Previous studies have also demonstrated a ‘modality-appropriateness’ effect in time perception in which time estimates are “pulled” toward the modality with the lower variance (van Wassenhove et al., 2008).

The specifics of Bayesian cue combination are that sensory estimates of time (*t*_*S*_) are drawn from a normal distribution:

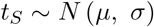

With a specific mean and variance, depending on the fidelity of the sensory modality conveying the temporal information. When two sensory estimates are available, each conveying a measure of duration, both are combined into a multisensory estimate. Assuming one signal is from a sensory (S) modality (e.g. auditory, visual) and the other is from movement (M), each with its own mean and variance, the combined sensorimotor (SM) estimate is given as:

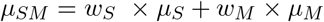

Wherein each independent channel is weighted. In this way the peak of the SM distribution will be closer to whichever sensory estimate has the greater weight. The weights themselves for each estimate are then calculated (for M, as example) as:

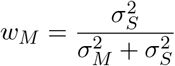

That is, the weight for a given estimate is the ratio of the other modality’s variance over the sum of the variances of both modalities. For the variance of the combined SM estimate, this can be expressed as the product of the other variances divided by their sum, or:

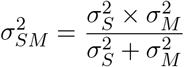

The essential result of this combination is that the SM estimate will always be more precise than either of the unisensory estimates, alone. By calculating these equations across a range of values, predictions can be made of the behavior of multisensory estimates across a wide range of possible variances (Figure 1D, E, F).

**Figure 1:**
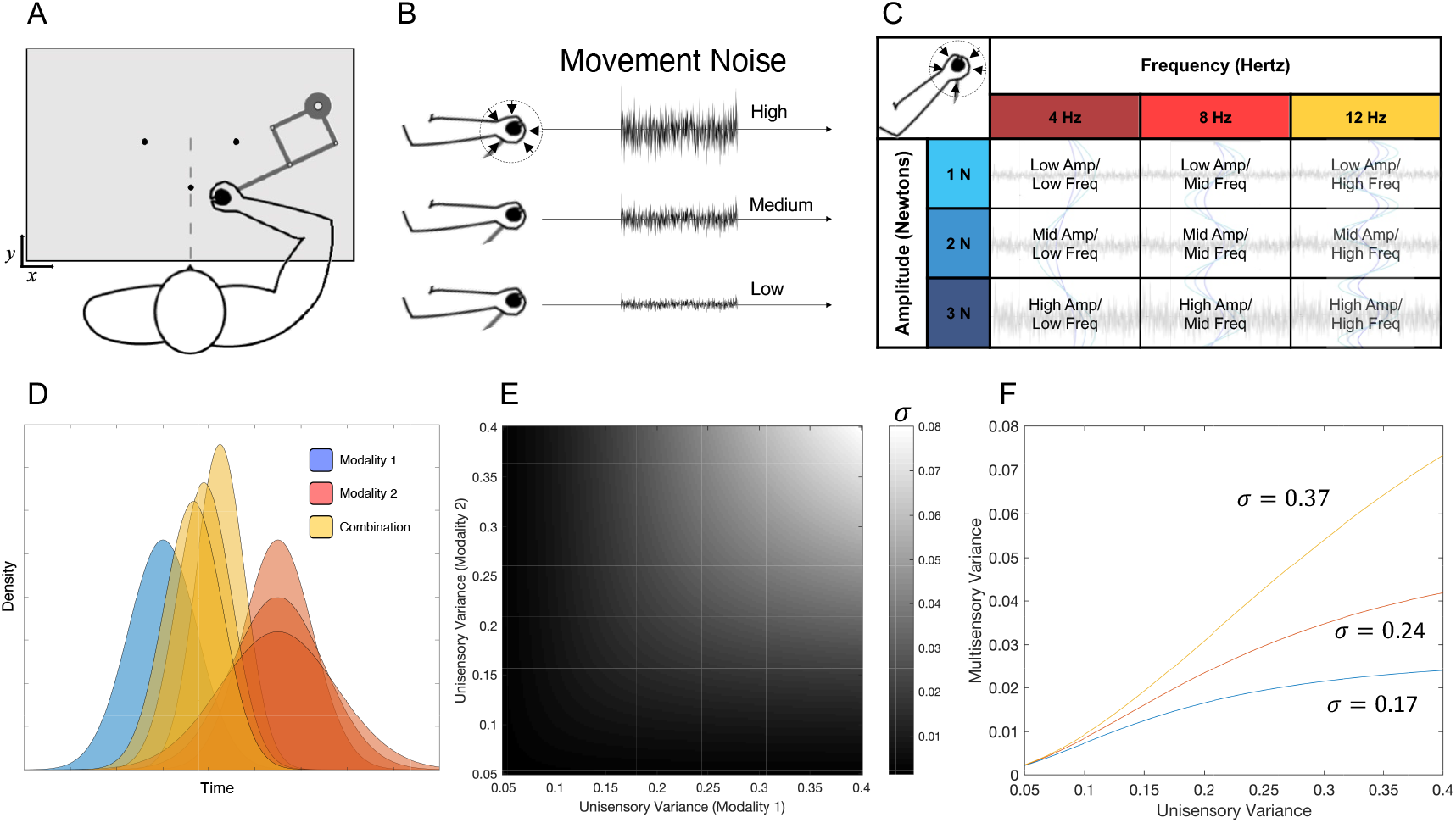
Experimental setup and Cue Combination Model. **A)** Subjects sat facing and held a robotic arm manipulandum that was kept below an occluding viewscreen. The experimental paradigm required subjects to begin each trial by moving the arm to a central starting location. On a given trial, subjects were required to categorize time intervals into “short” and “long” duration categories by moving the arm cursor to one of two response locations, counterbalanced between subjects. **B)** On any given trial, subjects experienced movement noise in the robotic arm, generated by introducing a tremor on the arm itself, in a random direction between trials, across one of three noise levels (low, medium, high). **C)** Movement noise was designed across three levels of two dimensions: the amplitude of the noise, expressed in newtons, or the frequency of the noise, expressed in frequency. **D)** Bayesian Cue Combination predictions, in which two signals from separate modalities are combined optimally to form a combined distribution that is more precise than either distribution alone. If one modality is made less precise (wider width), the combined distribution also lowers in precision, while becoming weighted progressively in location towards the more precise modality. **E)** Multisensory combination widths, expressed as a function of the precision of both unisensory widths, provide predictions of precision as a function of either increasing or decreasing levels of noise. **F)** When the precision of one modality is kept constant, increasing the variance of the other distribution generates variance curves that depend on the base precision of the first modality. Three levels of base precision are presented here (*σ*= 0.17, 0.24, 0.37); with less precise base modalities, increasing noise in the second modality has progressively larger effects.

Two predictions made by these equations are that 1) sensorimotor estimates of duration will be combined optimally, and 2) the sensorimotor estimate will depend on the precision of both modalities. Recently, we tested the first prediction in a study in which subjects measured either a sensory (auditory) time interval, or timed their own movements, or both. We found that subjects’ perception of either auditory tones or their own movements were indistinguishable in terms of their precision, yet each produced systematic errors, with subjects overestimating auditory tone intervals and underestimating the duration of their own movements. Notably, the combined estimate was in between both, and also the most accurate. However, despite this improvement, subjects exhibited a suboptimal combination of both modalities; yet, the degree of suboptimality depended on the individual’s overall level of precision, with more precise subjects closer to the optimal estimate (De Kock et al., 2023).

For the second prediction of the cue combination framework, when a more precise modality (i.e., movement) becomes less reliable, or “noisy”, its influence on the estimate will decrease; therefore, if movements are made unreliably or with uncertainty, they should have less of an influence than other modalities on time estimates ((De Kock et al., 2021)). Further, the size of this effect should vary with the baseline level of precision of the sensory modality. That is, if the sensory estimate of duration is already precise, then increasing noise in movements will have less of an effect with higher levels of noise. Conversely, if the sensory estimate is already noisy, increasing noise in movements will have a larger effect.

How can noise be introduced into motor movements? Critically, we conceive of a noisier movement as one that is less *reliable*. In other words, one where subjects feel their movements cannot be trusted or depended upon. A common example of motor noise is the experience of tremor, which in clinical cases (e.g., Parkinson’s Disease and essential tremor) can severely disrupt motor control (McAuley and Marsden, 2000). In the present study, we applied a tremor to healthy participants making arm movements to test whether this source of noise disrupted timing performance. Participants used a robotic arm to perform a temporal categorization task while experiencing variable tremor frequencies and amplitudes. We hypothesized that this source of motor noise would disrupt timing precision, but not accuracy, as predicted by the Bayesian cue combination framework.

## Results

### Experiment 1: Auditory Categorization

To begin, our first experiment tested 24 individuals on an auditory temporal categorization task (also referred to as temporal bisection), in a manner similar to our previous work (De Kock et al., 2021; Wiener et al., 2019). Specifically, human participants sat facing forward while holding a robotic arm manipulandum under an occluding viewscreen (Figure 1A). On a given trial, an auditory tone was played for one of seven possible intervals between one and four seconds (log-spaced). Subjects were required to move the cursor indicating arm position to one of two response locations, equidistant from the starting location on a given trial. Crucially, we introduced three levels of movement “noise” while subjects moved the robotic arm, expressed as a tremor applied to the handle (Figure 1B). We characterized noise across two different dimensions, in which both the amplitude of the tremor (1-3 Newtons) and its frequency (4-12Hz) could vary from trial to trial (Figure 1C). Our decision to parametrically vary tremors across these two dimensions was driven by our agnostic view to what the motor system may consider “noisiness”. Additionally, the direction of the tremor varied randomly between trials. Similar to our previous studies, subjects were required to categorize the interval as quickly, but as accurately as possible, and only enter the response location once the auditory tone had completed; further, subjects were required to maintain movement throughout the trial, with any violation leading to trial termination.

Analysis of choice and RT data also proceeded similarly to our previous reports, with choice data fit with a psychometric curve to calculate the bisection point (BP; 0.5 probability of choosing “long”) and the coefficient of variation (CV; half difference between 0.75 and 0.25 probability points divided by the BP); the BP reflects the level of bias in categorization, whereas the CV reflects the precision (Figure 2A). A repeated measures ANOVA revealed that there were no significant effects of amplitude (*F* _(2,46)_ = 0.05, *p* = .955) or frequency (*F* _(2,46)_ = 1.68, *p* = .197), nor was there an interaction effect of the two (*F* _(4,92)_ = 1.42, *p* = .233) on BP (Figure 2C). Therefore, participants were not distracted by the induced tremor and were able to accurately complete the task (Fig 2). There were also no significant effects of frequency (*F* _(2,46)_ = 1.37, *p* = .264), amplitude (*F* _(2,46)_ = 0.13, *p* = .879), nor an interaction effect of frequency and amplitude (*F* _(4,92)_ = 0.08, *p* = .988;) on reaction time (RT) (Figure 2D).

**Figure 2:**
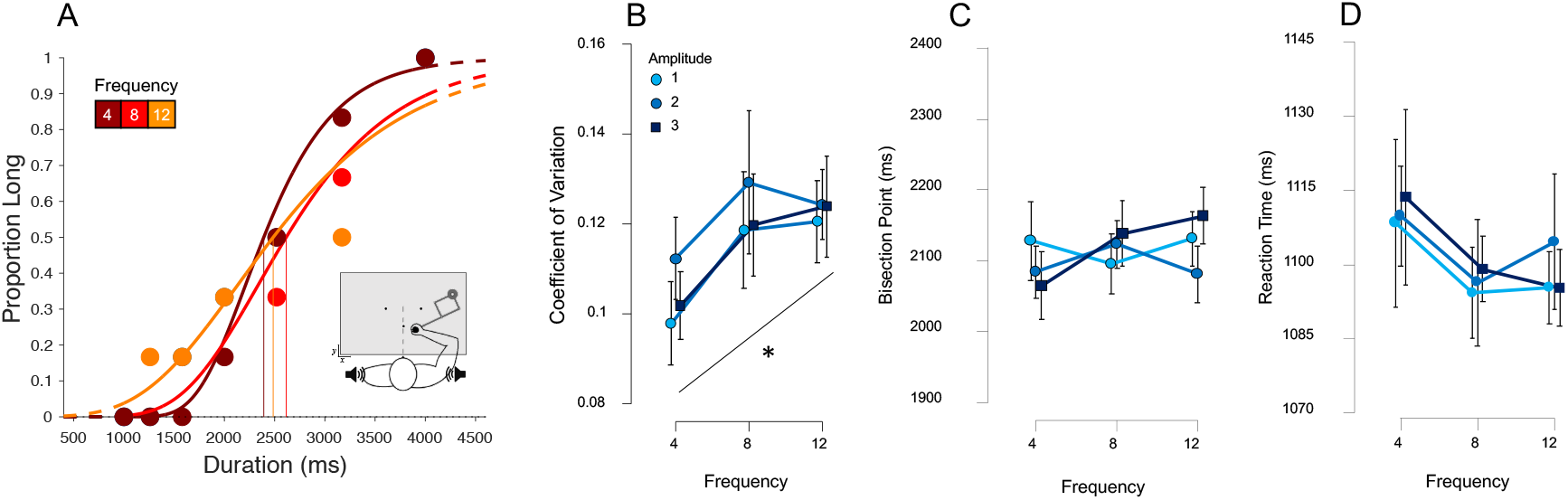
Results from Experiment 1 (Auditory Categorization). **A)** psychometric curves fit to response proportions for a representative subject exhibiting a decrease in precision with increasing motor noise frequency, collapsed across amplitude; vertical lines indicate the Bisection Point (0.5 probability of classifying ‘long’). **B)** Average coefficient of variation (CV) for all subjects across all noise amplitudes and frequencies, demonstrating an increase for frequency, with no change for different amplitudes. **C,D)** Average Bisection Points and reaction times across all conditions demonstrating no significant differences across either noise type. Error bars represent standard error; asterisk represents a linear effect at p*<*0.05.

However, a repeated measures ANOVA of the CV revealed a significant effect of frequency (*F* _(2, 46)_ = 3.85, *p* = 0.028, *η*^2^_p_ = 0.143); there was no significant effect of amplitude (*F* _(2,46)_ = 0.46, *p* = .636) nor an interaction effect of amplitude and frequency (*F* _(4,92)_ = 0.10, *p* = .983). A further examination revealed this to be a linear effect, with CV values generally increasing with frequency, (*t* = 2.425, *p* = .019, Cohen’s *D* = 0.715) suggesting precision decreased as frequency increased. (Figure 2B)

A control analysis on the direction of the movement tremor was also conducted, by dividing responses into three separate bins, depending on the direction of the tremor (bin 1: 0-60°; bin 2: 61-120°; bin 3: 121-180°). Here, no effect of tremor direction was detected for either the BP (*F* _(2,46)_=0.321, *p =* .727) or the CV (F_(2,46)_=0.755, *p* = .476), suggesting the direction of the tremor had no effect on either bias or precision.

In addition to the response variables, we also wanted to evaluate the parameters of movements made while encoding the duration, specifically movement length and force (supplementary figure 1). A repeated measures ANOVA for movement length revealed no significant effect of frequency (*F* _(2,46)_ = 2.93, *p* = .063) or amplitude (*F* _(2,46)_ = 0.01, *p* = .994), nor an interaction effect (*F* _(4,92)_ = 1.97, *p* = .106); Although the frequency effect is not significant, the data and our hypotheses suggest a linear effect of frequency but not amplitude on movement length; therefore, we explored the linear comparison contrast of frequency across all levels of amplitude which revealed a significant effect (*t* = -2.24, *p <* .05). As expected from previous research (Wiener et al., 2019; De Kock et al., 2021), movement length increased with duration (*F* _(6,138)_ = 254.07, *p <* .001).

A repeated measures ANOVA of movement force revealed a significant effect of frequency (*F* _(2,46)_ =34.41, *p <* .001), amplitude (*F* _(2,46)_ = 5.26, *p <* .05), and an interaction effect of frequency and amplitude (*F* _(4,92)_ = 9.44, *p <* .001). Post hoc analysis of frequency showed that movement force was not significantly higher at the lowest frequency (Freq4) compared to the mid frequency (Freq8; *t* = 2.36, *p* = .081) and significantly higher compared to the highest frequency (Freq12; *t* =8.17, *p <* .001). Movement force was also significantly higher at the mid frequency (Freq8) compared to the highest frequency (Freq12; *t* = 5.28, *p <* .001). Post hoc analysis of amplitude showed that movement force was significantly lower at the lowest amplitude (Amp1) compared to the mid amplitude (Amp2; *t* = -3.70, *p <* .01) and the highest amplitude (Amp3; *t* = -2.94, *p <* .001). Movement force was also significantly lower at the mid amplitude (Amp2) compared to the highest amplitude (Amp3; *t* = -4.97, *p <* .001).

### Computational Modeling

The results of Experiment 1 support, in principle, the predictions of the Bayesian Cue Combination model. That is, if two sensory modalities both convey estimates of time, these estimates will be combined optimally to improve perceived duration. However, if one of those modalities becomes noisier, and so less reliable, the overall precision will decrease as the noise increases (Hartcher-O’Brien et al., 2014). Further, the decrease in precision should diminish with increasing noise, depending on the precision of the unchanged modality. If movement represents a sensory channel for estimates of duration, then increasing noise in movements should decrease the precision of time estimates. This prediction was borne out in our data; however, alternative explanations to the findings exist. Indeed, decreases in timing precision (increases in CV) can be explained by numerous effects in the timekeeping process; noisier time estimates may result from a poorer ability to encode time, or from an impairment in remembering those durations, or also a difference in how those durations are judged (Allman et al., 2014). In our previous report, we had shown that increasing movement viscosity shifted response bias (changes in the BP), while sparing precision (De Kock et al., 2021). There, we found through computational modeling of behavior that the effect likely arose from a difference in perception, rather than a change in decision-making. We chose to take a similar approach here.

To begin, we opted to again to employ a drift-diffusion modeling (DDM) framework. The classic DDM is able to account for a wide variety of effects in choice and RT by accounting for the shape of response distributions across different experimental levels (Ratcliff, 1978). The typical DDM assumes that information is accumulated over time in a noisy stochastic process towards one of two response thresholds, with a given boundary separation. The rate of accumulation is determined by the drift rate parameter (v), and is corrupted by white noise on a moment-by-moment basis during the accumulation process, until the boundary is reached, given by the threshold parameter (a). Additional parameters include a delay in the initiation of accumulation, known as non-decision time (t) and starting-point bias towards one of the two boundaries (z).

Here, we used the HDDM (version 0.9) package for model construction and simulations (Wiecki et al., 2013). HDDM allows for the construction and fitting of hierarchical DDMs by constraining individual-level parameter estimates on the basis of group-level ones in addition to prior distributions for the given parameters. For the present study, we employed the recently updated “Likelihood Approximation Network” (LAN) extension to HDDM, in which a wider number of possible models are now supported by training through artificial neural networks (Fengler et al., 2021). These model extensions include those with collapsing boundary or leak parameters. Of relevance to the present study, the LAN extension also includes a so-called Lévy-Flight model (Voss et al., 2019). In this model, the noise for the momentary sensory evidence is modified by the parameter (alpha), which determines the shape of the noise distribution. This parameter, which ranges between 1 and 2, interpolates the shape of the noise distribution between a Gaussian distribution at higher values and a Cauchy distribution at lower values. The Cauchy distribution includes heavy tails in both directions, and so allows for large “jumps” in evidence accumulation towards either of the decision boundaries. As the Cauchy is narrower overall than the Gaussian, the accumulation process may be less noisy moment-to-moment, yet overall may shift randomly by a large amount.

For our model simulations, we suggest that “noise” may be considered by changing either the drift rate, threshold, or alpha parameters (Figure 3A). To accomplish this, we began by fitting a DDM-Lévy model to our full dataset, with all parameters included [v, a, t, z, alpha], but only duration as a conditional parameter. We chose to include these parameters on the basis of our previous work and others demonstrating these parameters (excluding alpha) as the best for accounting for behavior on this particular task, and also to follow a-priori assumptions. Once the fitted parameters were obtained, we next simulated three separate datasets using these same parameters, but by varying each of the three parameters described above across three different levels (see Methods).

**Figure 3:**
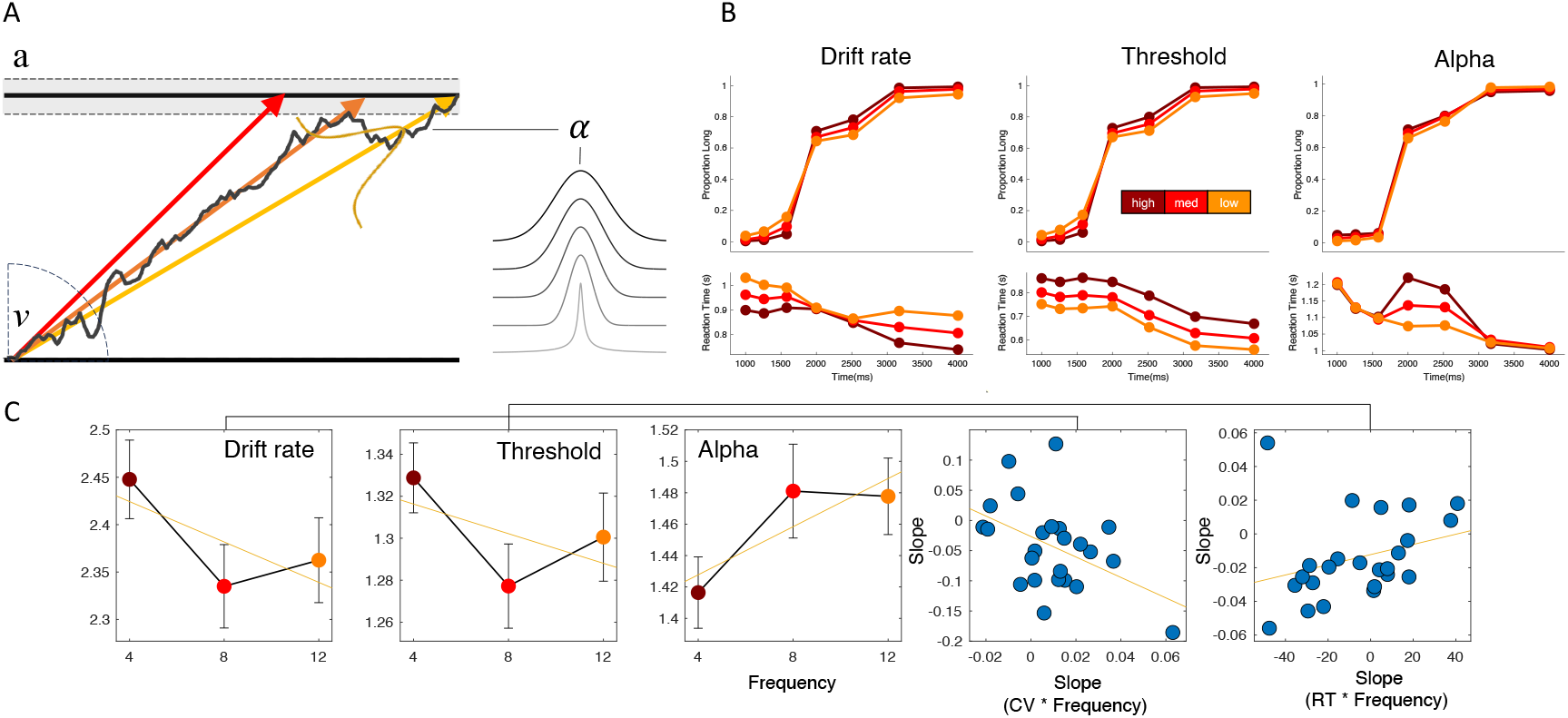
Computational modeling using a Drift Diffusion Model framework with Levy Flights. **A)** DDM framework in which evidence is accumulated to a boundary for categorizing a time interval as long or short. The rate of accumulation on a given trial is determined by the drift parameter (v; colored lines represent three possible drift rates on different trials), while the boundary for the response is determined by the threshold parameter (a; dashed lines represent different possible levels on different trials); additionally, the momentary noise of the distribution is determined by the noise transformation parameter (alpha) which determines the relative shape of momentary noise, interpolating between a Gaussian distribution and a Cauchy distribution (heavy tailed) at higher and lower levels, respectively. **B)** Choice and RT findings from simulated models with each of the three possible parameters varying over three different levels. All three models lead to changes in choice precision by increasing parameter values, but with different RT predictions. **C)** Fits of the DDM-Levy model, collapsed across amplitude, reveal significant changes in all three parameters with increasing movement noise frequency (left 3 panels). However, only the drift rate effect correlates with the CV effect of behavior, whereas the threshold effect correlates with changes in RT.

For the drift rate, when considered as an absolute value (that is, the drift rate may be considered either signed, pointing towards one boundary or another, or unsigned, indicating overall slope) lower drift rates would indicate a slower accumulation process, which has been shown to relate to the overall signal-to-noise ratio in perception. We simulated three separate datasets with three drift rates (high, medium, low) and observed that as the drift rate lowers, the psychometric curves becomes less steep, indicating a decrease in precision. Notably, the RT shape across intervals also changes, with lower drift rates associated with longer RTs, but only for the longest and shortest interval in the stimulus set. For the threshold parameter, lowering the threshold also resulted in a decrease in precision, yet here the RT distribution shifted uniformly across all intervals, with responses becoming quicker for lower threshold. This effect is due to lower thresholds leading to faster responses, which require less evidence accumulation. For the alpha parameter, we again observed a decrease in precision, but for lower values of alpha instead of higher values. That is, as the noise more closely approximated a Cauchy distribution, the shallower the psychometric curve became. Further, RTs increased with lower alpha values, but only for the middle durations; this effect likely results from a longer amount of time necessary with a more constrained evidence accumulation regime to reach a specified boundary, thus lengthening the time before a response is committed (Figure 3B).

Altogether, all three models could provide an explanation for decreases in precision resulting from increases in noise, yet with differing predictions for RT. After fitting the full model to our data, we observed that all three parameters additionally shifted with changes in Frequency (Figure 3C). More specifically, we observed a decrease in drift rate as frequency increases (*F* _(2,46)_ = 9.962, *p <* 0.001), in addition to a decrease in threshold (*F* _(2,46)_ = 14.661, *p <* 0.001), and an increase in alpha (*F* _(2,46)_ = 4.98, *p =* .011). Across these parameters, we note that only the change in drift rate and threshold were consistent with the model simulation results; an increase in alpha predicts better precision, rather than a decrease as observed in our behavioral data. However, for the drift rate and threshold, either may match the behavioral data. To determine which, we calculated the slope of a linear regression across frequency for both drift rate and threshold, and correlated those values between-subject with the slope of the CV values across frequency. Here, we observed that only the drift rate effect significantly correlated with the CV effect [Pearson’s *r* = -0.451, *p* = .026; Spearman’s *r* = -0.413, *p* = .044], whereas the threshold effect did not [Pearson’s *r* = -0.213, *p* = .317; Spearman’s *r* = -0.281, *p* = .182]. Conversely, we found that the threshold effect could explain changes in RT across frequency levels [Pearson’s *r* = 0.296, *p* = .159; Spearman’s *r* = 0.413, *p* = .044; we note the lack of a Pearson effect here is likely driven by an outlier, for which the Spearman effect is not affected], whereas drift could not [Pearson’s *r* = -0165, *p* = .438; Spearman’s *r* = -0.086, *p* = .685]. Although the RT effect was not significant, we note that theoretically the threshold parameter should be able to explain any between-subject differences in RT.

### Experiment 2: Visual

The overall findings of Experiment 1 and computational modeling both support the notion that increasing the frequency of movement noise leads to a decrease in perceptual timing precision in accordance with Bayesian cue combination. However, recall in the cue combination framework that the overall effect of increasing noise on one modality will depend on the base level of precision in the other modality; if the unchanged modality’s precision is high, then the impact of increasing noise in the second modality will diminish with higher levels of noise, whereas if the unchanged modality’s precision is low, then increasing noise in the second modality will have a larger effect with less diminishment. To test this possibility, we repeated our temporal categorization task in a new sample of subjects (n=24), but with the visual modality used for timing instead of auditory. It is well documented that the fidelity of perceptual timing is worse for visual than for auditory stimuli, such that precision is lower for the former than the latter (Wiener et al., 2014; Shi et al., 2013; van Wassenhove et al., 2008). In place of an auditory tone, the visual interval was demarcated by a global change in luminance of the viewscreen (see Methods). This was done so that subjects would be able to easily attend to the onset/offset of the stimulus regardless of where they were looking on the screen or where the cursor was located.

As in Experiment 1, we also analyzed BP and CV as measure of bias and precision (Figure 4A), respectively, using repeated-measure ANOVAs. Similar to those results, there was no effect of amplitude (*F* _(2,46)_ = 2.23, *p* = .119) or frequency (*F* _(2,46)_ = 0.42, *p* = .66), nor an interaction effect (*F* _(4,92)_ = 0.39, *p=* .814) on BP, again suggesting participants were also not distracted by the noise or stimuli and were able to accurately complete the task (Figure 4C). There was again no effect of frequency (*F* _(2,46)_ = 2.28, *p* = .114), amplitude (*F* _(2,46)_ = 0.83, *p* = .443), or an interaction effect (*F* _(4,92)_ = 0.82, *p* = .515) on reaction time (RT) (Figure 4D). A control analysis on tremor direction again was conducted on choice responses, which once again failed to find any effect on either the BP (*F* _(2,46)_ = .067, *p =* .935) or the CV (*F* _(2,46)_ = 0.455, *p =* .637).

**Figure 4:**
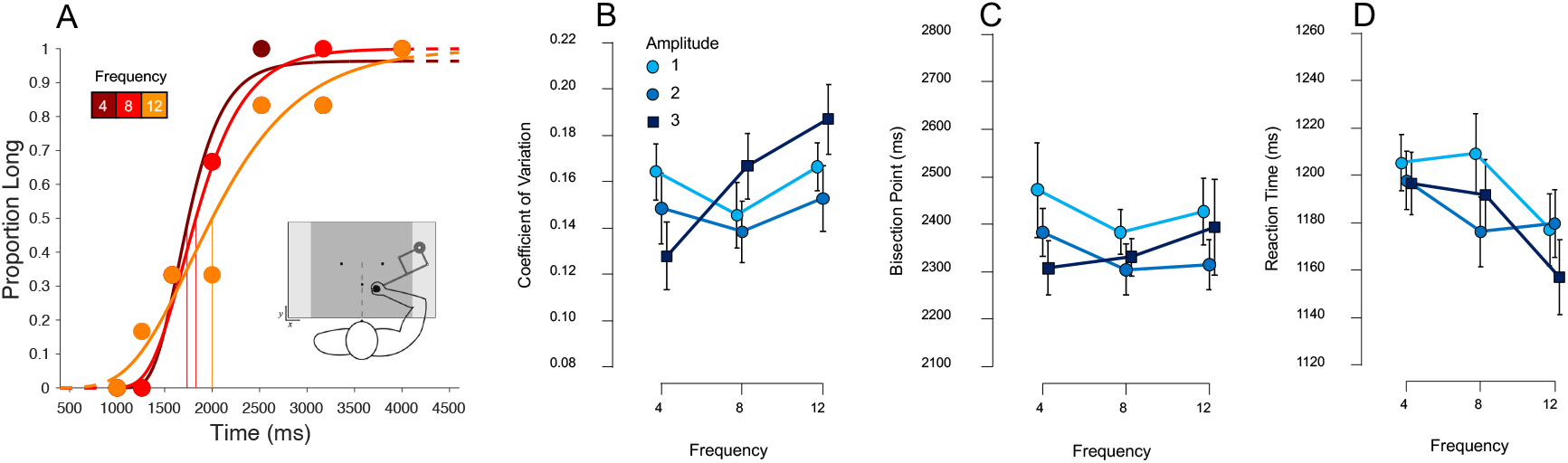
Results from Experiment 2 (Visual Categorization). **A)** Psychometric curves fit to response proportions for a representative subject again exhibiting a decrease in precision with increasing movement noise frequency (highest amplitude). **B)** Average CV for all subjects across all noise amplitudes and frequencies; in contrast to Experiment 1, an increase in CV across frequency was observed only for the highest amplitude noise (3 Newtons). **C,D)** Average Bisection Points and reaction times across all conditions again demonstrating no significant differences across either noise type. Error bars represent standard error.

For the CV we again found no effect of amplitude (*F* _(2,46)_ = 1.01, *p* = .374) or an interaction effect of amplitude and frequency (*F* _(4,92)_ = 1.58, *p* = .188). However, we did not observe a main effect of frequency (*F* _(2,46)_ = 2.61, *p* = .084, *η*^2^_p_ = 0.102); we note that the linear contrast was significant (*t* _(46)_=2.121, *p* = .039, *D* = 0.625), indicating an overall increase in CV with higher frequencies. Following our a-priori hypotheses, we analyzed Simple Main Effects with frequency as the Simple Effect Factor and amplitude as the Moderator. Here, we found a significant effect of frequency only for the highest amplitude of three Newtons (*F* _(2,46)_ = 3.844, *p* = .029), but not for either of the lower amplitudes (*p* = .414 and .766, respectively), indicating that the CV did significantly increase, but only for the highest amplitude (Figure 4B).

We again analyzed movement length and force in addition to the response variables (supplementary figure 1). A repeated measures ANOVA of movement length showed no significant effect of amplitude (*F* _(2,46)_ = 2.68, *p* = .079) and no significant effect of frequency (*F* _(2,46)_ = 2.06, *p* = .139) nor an interaction effect (*F* _(4,92)_ = 0.96, *p* = .431). Given the marginal effect of amplitude and the linear nature of the data we decided to explore the linear comparison of amplitude averaged over all levels of frequency which revealed a significant effect (*t* _(46)_ = 2.30, *p <* .05). As expected from previous research (Wiener et al., 2019; De Kock et al., 2021) and Experiment 1 results, movement length increased with duration (*F* _(6,138)_ =309.25, *p <* .001).

A repeated measures ANOVA on movement force, however, revealed a significant effect of amplitude (*F* _(2,46)_ = 4.04, *p <* .001), frequency (*F* _(2,46)_ = 43.20, *p <* .001), and an interaction effect (*F* _(4,92)_ = 4.76, *p <* .001). Post hoc analysis of amplitude showed that movement force was significantly lower at the lowest amplitude (Amp1) compared to the mid amplitude (Amp2; *t* = -3.40, *p <* .01) and highest amplitude (Amp3; *t* = -5.50, *p <* .001). It was also significantly lower at the mid amplitude (Amp2) compared to the highest amplitude (Amp3; *t* = -4.97, *p <* .001). Post hoc analysis of frequency showed that movement force was significantly higher at the lowest frequency (Freq4) compared to the highest frequency (Freq12; *t* = 7.24, *p <* .001) and not significantly higher than the mid frequency (Freq8; *t* = 7.83, *p* = .081). Movement force was also significantly higher at the mid frequency (Freq8) compared to the highest frequency (Freq12; *t* = 7.83, *p <* .001).

### Cross-Modal Comparisons

In order to investigate differences between the two modalities (Exp 1: Auditory, Exp 2: Visual), we conducted an mixed-model ANOVA with modality as the between-subjects factor. Although at first glance it appears that movement force was overall higher in the visual modality (Exp. 1) than in the auditory modality (Exp. 2), this difference was not significant (*F* _(1,46)_ = 0.92, *p* = .341). Movement length was found to increase as frequency increased in the auditory modality (Exp. 1); however, in the visual modality (Exp. 2) movement length increased as amplitude increased. We again compared across modalities and found that movement length was not significantly longer in the visual modality (Exp. 2) compared to the auditory modality (Exp. 1; *F* _(1,46)_ = 3.47, *p* = .069). Together these findings suggest that increases in the size of the tremor led to increased movement force and longer movement lengths whereas increases in the speed of the tremor led to decreased movement force and so shorter movement lengths. Previous research has suggested that increases in movement length lead to increases in time perception (Wiener et al., 2019; De Kock et al., 2021); therefore, a logical next step was to compare the BP across the two modalities to see if this was the case here as well. However, cross-modal analysis revealed no significant differences on BP between groups (*F* _(1,46)_ = 2.83, *p* = .099). In addition to response variables and movement parameters, we also verified that there were no significant differences of modality on RT (*F* _(1,46)_ = 1.48, *p* = .230). This finding further supports the suggestion that the effects of frequency and amplitude on the different movement parameters did not influence the perception of time but was simply due to the nature of controlling the robotic arm with increasing size and speed of movements.

Lastly, and most important, was the cross-modal comparison of CV which revealed precision was overall lower (higher CV values) in the visual modality compared to the auditory modality (*F* _(1,46)_ = 10.01, *p <* .01, *η*^2^_p_ = 0.179). Additionally, we evaluated the size of the effect of frequency on each modality as, according to the cue combination framework, the influence of noise on modality should depend on the base precision of the unchanging modality, and so if the visual modality is less precise, then noise should have a larger effect. To test this, we calculated slope values for each effect and compared them. For the auditory group, we used the average effect of frequency, collapsed across amplitude, whereas for the visual group, we only used the effect of frequency for the highest amplitude (Figure 5). Here, a significant difference was found, with the effect of noise having a larger influence for the visual group than the auditory one [Mann Whitney U = 170, *p =* .014, rank biserial correlation = -0.41].

**Figure 5:**
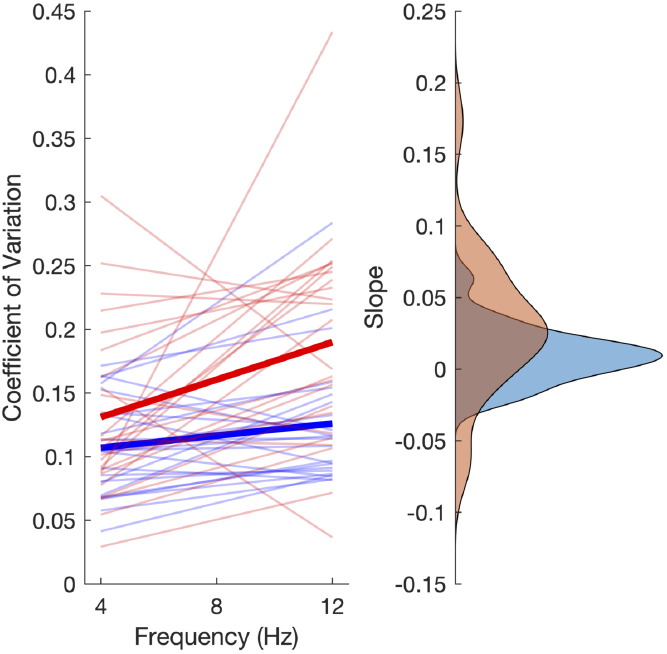
Slope comparisons between experimental groups. Left panel displays linear regressions of the coefficient of variation for individual subjects and group averages (bold lines; red = visual; blue = auditory) across the three noise frequencies. Right panel displays density estimates of the distribution of slope values for each. We observed that subjects performing the task with visual stimuli, in addition to higher CV values overall, exhibited a larger influence of noise frequency, as exhibited by a significantly higher slope value for these subjects.

## Discussion

The overall purpose of these experiments was to test the proposed Bayesian cue combination framework which suggests that movement serves as an additional channel of temporal information that has high precision and low temporal fidelity which improves the precision of time perception. We therefore considered, if movement became unreliable or noisy, we should see a decrease in the precision of timing in that, it will be pulled toward the more precise sensory input. We additionally wanted to investigate whether this effect would differ between auditory and visual stimulus. Given previous findings that auditory temporal perception is more precise than visual (Mioni et al., 2016; Wiener et al., 2014), we hypothesized that noisy movements would lead to less precise estimations of time with visual stimuli compared to auditory.

Different levels of amplitude or frequency did not have a significant effect on accuracy in either the auditory or visual version of the study suggesting that participants were able to appropriately complete the task and the noise did not serve as a distractor from the timing task. This distinction is important, as one might expect that increased tremors would lead subjects to pay less attention to the timing task. However, in that case, the prediction is that the time estimates themselves would also become more biased to be shorter (Fortin, 2003; Brown, 1997), which was not the case. There were also no significant effects of frequency or amplitude on RT within or between either experiment.

We did find an effect of precision in that, an increase in frequency but not amplitude led to a less precise perception of time for auditory stimuli whereas for visual stimuli the effect of frequency was only found at the highest level of amplitude. Specifically, for Experiment 1 (auditory) participants were most precise at the lowest frequency level (4 Hz) with a leveling off between the mid (8 Hz) and high frequency (12 Hz). For Experiment 2 (visual) we found the same pattern but only for the highest amplitude (3N), whereas for low (1N) and mid (2N) amplitudes there were no significant differences in precision. Therefore, when participants timed auditory tones, increasing the speed of the tremor but not the size of the tremor caused a decrease in precision. One explanation for this difference is the difference in baseline precision between auditory and visual modalities; that is, since auditory time estimates are already very precise, small disruptions to movements will lead to a change in their overall precision. In contrast, since visual time estimates are already less precise, a larger amount of noise is necessary in movements to induce an effect, as the noise in movements must exceed the noise of visual time estimates such that the cue combination begins to favor them over movements. As such, larger noise is only achieved in the visual experiment at a high amplitude.

The computational modeling conducted in our study allowed us to identify possible sources of noise in the perceptual process. By relying on a DDM framework that incorporates Lévy Flights, we observed that the behavioral effect could be explained by a shallower drift rate. Decreases in drift rate have been associated with a lower signal-to-noise ratio, and so relate to the rate of evidence accumulation in the perceptual process (Voss et al., 2004; Palmer et al., 2005; Rohenkohl et al., 2012). In our case, when the drift rate was lower, we observed a similar finding to our behavioral data. Further, fits of this model to the behavioral data revealed that changes in drift rate correlated with changes in precision, supporting the conclusion that the effects were driven by perception-level changes, rather than biases in decision-making or changes in strategy.

As for movement parameters during temporal encoding, in the auditory version (Exp. 1) we found there to be a linear effect of frequency but not amplitude on movement length; specifically, movement length significantly decreased linearly with an increase in frequency for the two higher amplitude levels (Amp2 and Amp3) but not at the lowest amplitude (Amp1). Notably, for the visual version of the study (Exp. 2) there was a linear effect of amplitude but not frequency where movement length significantly increased linearly with an increase in amplitude for the lowest frequency (Freq4).

We also observed differences in movement length between experiments in that movement length was overall longer in the visual study (Exp. 2) compared to the auditory study (Exp. 1); however, it was only marginally significantly different. Given previous findings and in line with proposed framework, we would expect a difference in movement length would lead to a biasing effect of time perception; however, there was no a significant difference between the two experiments on BP which suggests that movement length did not influence time perception in this case.

The results of our experiment have particular implications for the study of movement disorders. Notably, time perception abilities are impaired in pathologies of movement, such as Parkinson’s Disease (Singh et al., 2021), Huntington’s Disease (Lemoine et al., 2021), or Cerebellar Degeneration (Breska and Ivry, 2021), but also in Essential Tremor (Pedrosa et al., 2016), Tourette’s Syndrome (Vicario et al., 2010), and Dystonia (Conte et al., 2017). Conversely, individuals with highly trained coordination, such as professional athletes or musicians, exhibit enhanced timing abilities (Cicchini et al., 2012; Chen et al., 2016). Our observation that introducing a tremor to otherwise healthy individuals disrupts perception suggests an intrinsic link between motor symptomatology and perceptual processes. This link may further go beyond perception into the cognitive domain as well. Indeed, work with subjects with Attention Deficit Hyperactivity Disorder (ADHD), where well-known timing disruptions exist (Smith et al., 2002), has shown that ancillary movements can lead to improvements in perceptual processes (Hartanto et al., 2016). Similarly, recent work in Parkinson’s patients has linked deficits in sensorimotor timing to cognitive impairments (Singh et al., 2021). A corollary, implied by the present findings, is that by improving motor symptoms one could also improve both perception and cognition. Therefore, one might suggest that motor rehabilitation can also lead to other benefits in these patients. In the case of timing, it is possible to improve sensorimotor estimates through repeated training, which can lead to functional and morphological changes in sensorimotor brain regions (Bueti et al., 2012), yet whether this also improves other symptoms is unknown.

A second future avenue of research relates to the frequencies employed in the present study. Here, we chose the specific range (4-12Hz) to reflect that observed in motor system tremors (McAuley and Marsden, 2000). We note, however, that the higher end of the tremors is closer to that identified as the so-called sensorimotor “mu” rhythm (8-13Hz) (Pineda, 2005). While traditionally involved in movements, recent work has linked mu oscillations to motor-related changes in time-keeping processes, both for single intervals and rhythmic ones (Ross et al., 2022; Iwasaki et al., 2018). Notably, mu rhythms exhibit suppression during action initiation. One possibility, then, is the effects observed in the present study relate to a kind of resonance with mu oscillations via the induced tremor, thus leading to the observed disruption. Neural recordings, combined with a wider array of tremor frequencies, could shed light into this possibility, which would provide a distinct mechanism by which tremors exert their influence on the motor system.

Overall, our method, which relies on a novel use of a robotic arm to mimic tremors, was able to effectively alter timing performance. We suggest that our findings are not due to changes in attention or decision-making, but instead result from a fundamental change in perceptual processing, which follows from the Bayesian cue combination account. In conjunction with our other findings, we now find converging evidence to support the cue combination account for how movements influence perceptual time estimates, which in turn supports the hypothesis that movements *themselves* act as a timekeeping process with high fidelity.

## Materials and Methods

### Participants

A total of 48 participants that were right-hand dominant with normal or corrected-to-normal vision completed the following two experiments (Experiment 1: 24 participants - 15 female, 9 male, mean age = 23 years old; Experiment 2: 24 participants - 18 female, 6 male, mean age = 21 years old) from the University of California Davis student population and surrounding area. All participants were screened for handedness using the Edinburgh Handedness Survey (Oldfield, 1971) and provided consent as approved by UC Davis Institutional Review Board.

### Apparatus

Participants completed both experiments using a robotic manipulandum (KINARM End-Point Lab, BKIN Technologies). They were seated in an adjustable chair in front of the manipulandum at the height at which their forehead could rest comfortably on the apparatus’ headrest. The horizontal display was mirrored from the downward facing LCD monitor positioned above which occluded the participant’s view of most of their arm in order to reduce feedback of arm and hand position. Participants gripped the right handle of the apparatus and made movements within the screens perimeter and reaching movements to circular targets 0.5 cm in diameter, placed 14 cm apart on the sagittal axis of the body. Each target represented short and long responses counterbalanced across participants. The manipulandum continuously measured handle position, velocity, and force applied at a sampling rate of 1000 Hz.

### Procedures

#### Experiment 1: Auditory

Participants completed a temporal bisection task in which participants would move to the central target location where the manipulandum would lock in place. After 1000 ms, the warm-up phase would begin by the handle being released and the words ‘Get Ready’ being displayed on the screen. At this time, participants were told to move freely within the perimeter of the workspace. While participants moved, a tremor was mechanically induced by the manipulandum at one of three amplitudes (1, 2, 3 N) and frequencies (4, 8, 12 Hz), the direction of which was randomly selected from 180-degrees on each trial (note that the tremor rattled in both directions). Following a 2000ms delay, a 440 Hz tone sounded for one of seven durations between 1000-4000 ms (1000, 1260, 1580, 2000, 2520, 3170, 4000 ms). When the tone stopped sounding, the participants were to move to one of two response circles as quickly and accurately as possible to categorize the auditory tone as a short or long duration compared to all tones experienced so far (reference-free categorization). If a response was made prior to the auditory tone ending, or if subjects stopped moving, the trial was discarded and they were required to re-do the trial. The two targets were located at 105- and 75-degrees equidistant from the starting location with response assignment (short or long) counterbalanced between participants. A total of 378 trials were run per session.

#### Experiment 2: Visual

The goal of Experiment 2 was to test the impact of induced tremor on visual interval timing. The task parameters were identical to the auditory task, except that participants were required to categorize visual intervals presented as a change in the background luminance of the manipulandum display screen from black to gray (RGB values of 64,64,64, Hue 160, Luminance 60). After the screen reverted to black, subjects indicated whether the durations were ‘short’ or ‘long by reaching to the appropriate target. We tested visual intervals using this global background change rather than a fixation target to avoid spatiotemporal processing effects unrelated to duration encoding. A total of 378 trials again were run per session.

### Analysis

In Experiments 1 and 2, movement distance and force measures were taken for each trial. Movement distance was defined as the summed distance traveled (point-by-point Euclidean distance between each millisecond time frame) during the stimulus tone. Force was similarly defined as the summed instantaneous force during the stimulus tone. In Experiment 1, RT was defined as the time elapsed between tone offset and reaching one of the two choice targets, whereas in Experiment 2 it was the time between luminance offset and target. Outlier trials were excluded for RT values greater than three standard deviations away from the mean of a participant’s log-transformed RT distribution (Ratcliff, 1993). Additionally, we removed all trials with RTs below 200ms or above 2000ms, in order to avoid issues with model fitting. For each participant we plotted duration by average proportion of ‘long’ responses. From here, we used the psignifit 4.0 software package to estimate individual BP and coefficients of variation (CV) for all frequency and amplitude values (Schütt et al., 2016); all curves were fit with a cumulative Gumbel distribution to account for the log-spaced nature of tested intervals (Wiener et al., 2019; De Kock et al., 2021). The BP was defined as the 0.5 probability point on the psychometric function for categorizing intervals as ‘long’; the CV was defined as half the difference between 0.75 and 0.25 points on the function divided by the BP.

### Computational Modeling

To better dissect the results of Experiment 1, we decomposed choice and RT data using a drift diffusion model (DDM; (Ratcliff, 1978; Wiecki et al., 2013; De Kock et al., 2021). Due to the low number of trials available per condition, we opted to use hierarchical DDM (HDDM) as employed by the HDDM package (version 0.9.8) for Python (https://github.com/hddm-devs/hddm). In this package, individual subjects are pooled into a single aggregate, which is used to derive fitted parameters by repetitive sampling from a hypothetical posterior distribution via Markov Chain Monte Carlo (MCMC) sampling. From here, the mean overall parameters are used to constrain estimates of individual-subject estimates. HDDM has been demonstrated as effective as recovering parameters from experiments with a low number of trials (Wiecki et al., 2013).

A recent extension to the HDDM package, the Likelihood Approximation Network (LAN) module, allows for the “base” DDM to accommodate a wider variety of models (Fengler et al., 2021). Traditionally, the DDM consists of four parameters: the threshold difference for evidence accumulation (a), the drift rate towards each boundary (v), the starting point, or bias toward a particular boundary (z), and the non-decision time (t), accounting for remaining variance due to non-specific processes (e.g. perceptual, motor latencies). With the LAN module, additional parameters can be accessed and adjusted in model construction. For our purposes, we chose the so-called “Lévy-Flight” model extension, in which the noise in momentary evidence accumulation is modified by a parameter (alpha) which interpolates between a Gaussian distribution and a Cauchy distribution. Recent work has shown that the Lévy Flight model can accommodate many instances of 2-choice decision-making, with the additional feature that it can account for randomness in choice (Voss et al., 2019). We chose to use this model here, as the alpha parameter allows for another possible source of noise in the perceptual process.

Our initial model construction began by fitting the data from Experiment 1 using a so-called “full” DDM, as done in our previous work (De Kock et al., 2021). In this model, the only condition by which parameters vary is the duration presented on each trial; all parameters were set to vary. We chose this model setup to replicate both our previous work with this model, as well as other work on time categorization tasks demonstrating. Model construction was conducted using the HDDMnnStimCoding class. Model sampling was conducted using 10,000 MCMC samples, with a burn-in of 1000 samples and a thinning (retention) of every 5th sample. Individual model fits were assessed by visual inspection of the chains and the MC err statistic; all chains exhibited low autocorrelation levels and symmetrical traces. The resulting model contains seven values for each of the four parameters, reflecting each of the seven durations tested.

Once the model fits were obtained for each duration, we next proceeded to model simulation, so as to demonstrate how changing our three hypothetical parameters (v, a, alpha) could influence precision and RT. To do this, we used the simulator stimcoding class to generate data (1400000 trials each) from three separate models, with three levels within each model. The levels for each model were conducted by taking the parameter of interest for that model (v, a, or alpha) and multiplying them by 0.75 for each level; for example, if the drift rate for a given duration was 2, the next level would be 1.5, and the next would be 1.125.

Fits to the behavioral data to assess the effect of frequency and amplitude were conducted by creating a model in which all five parameters (v, a, t, z, alpha) varied according to the three levels of each (1,2,3 N for amplitude; 4, 8, 12 Hz for frequency). Model fitting used the same sampling method as described above, and again chain stability was assessed. We note that we attempted a similar modeling approach for Experiment 2, but found large instability and autocorrelations in the chains. This likely reflects the larger variability in Experiment 2 across all conditions.

## Supplementary Material

**Figure 6: Supplementary Figure 1.**
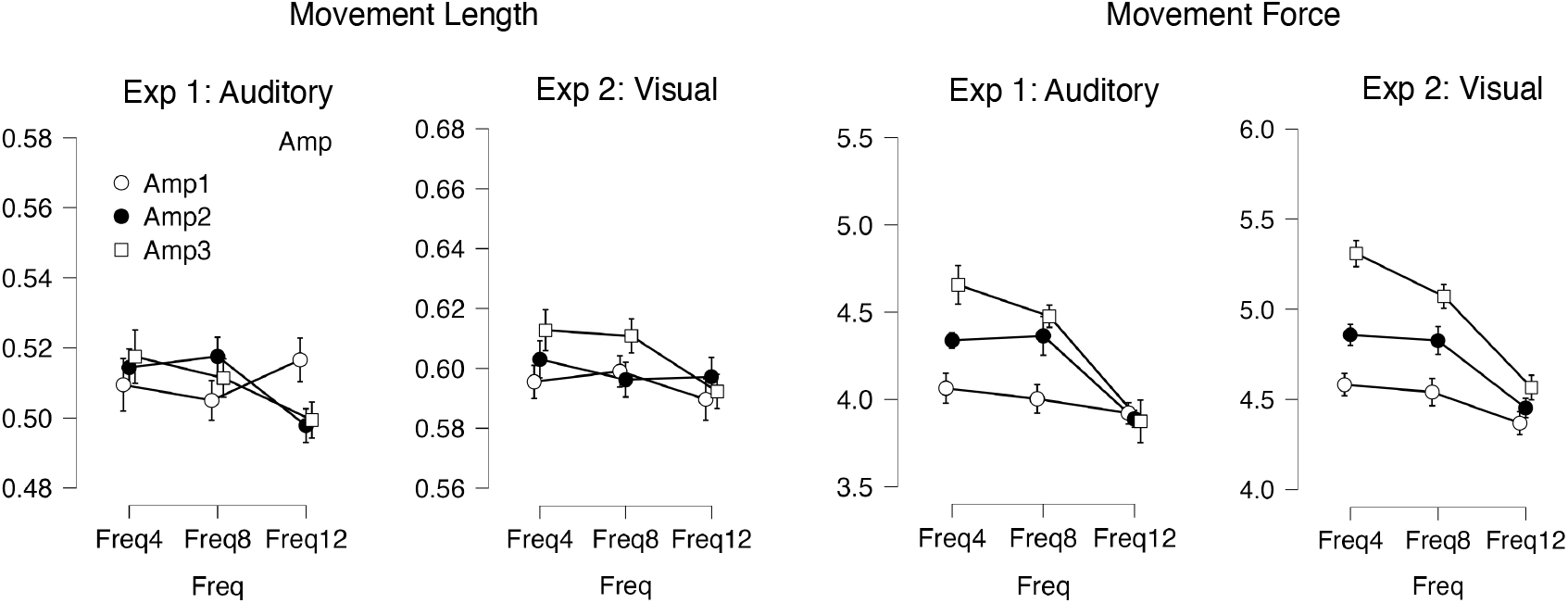
Movement length and force effects across both experiments.

## References

Alais, D., Burr, D., 2019. Cue Combination Within a Bayesian Framework, in: Multisensory Processes. Springer International Publishing, pp. 9–31. https://doi.org/10.1007/978-3-030-10461-0_2

Allman, M.J., Teki, S., Griffiths, T.D., Meck, W.H., 2014. Properties of the internal clock: first- and second-order principles of subjective time. Annu Rev Psychol 65, 743–71.

Brenner, E., van Dam, M., Berkhout, S., Smeets, J.B.J., 2012. Timing the moment of impact in fast human movements. Acta Psychologica 141, 104–111. https://doi.org/10.1016/j.actpsy.2012.07.002

Breska, A., Ivry, R.B., 2021. The human cerebellum is essential for modulating perceptual sensitivity based on temporal expectations. Elife 10.

Brown, S.W., 1997. Attentional resources in timing: interference effects in concurrent temporal and nontemporal working memory tasks. Percept Psychophys 59, 1118–40.

Bueti, D., Lasaponara, S., Cercignani, M., Macaluso, E., 2012. Learning about time: plastic changes and interindividual brain differences. Neuron 75, 725–37.

Chen, Y.H., Verdinelli, I., Cesari, P., 2016. Elite Athletes Refine Their Internal Clocks: A Bayesian Analysis. Motor Control 20, 255–65.

Cicchini, G.M., Arrighi, R., Cecchetti, L., Giusti, M., Burr, D.C., 2012. Optimal encoding of interval timing in expert percussionists. J Neurosci 32, 1056–60.

Conte, A., McGovern, E.M., Narasimham, S., Beck, R., Killian, O., O’Riordan, S., Reilly, R.B., Hutchinson, M., 2017. Temporal Discrimination: Mechanisms and Relevance to Adult-Onset Dystonia. Front Neurol 8, 625.

De Kock, R., Gladhill, K.A., Ali, M.N., Joiner, W.M., Wiener, M., 2021. How movements shape the perception of time. Trends Cogn Sci 25, 950–963.

De Kock, R., Zhou, W., Datta, P., Mychal, J.W., Wiener, M., 2023. The role of consciously timed movements in shaping and improving auditory timing. Proc Biol Sci 290, 20222060.

De Kock, R., Zhou, W., Joiner, W.M., Wiener, M., 2021. Slowing the body slows down time perception. Elife 10.

Doumas, M., Wing, A.M., Wood, K., 2008. Interval timing and trajectory in unequal amplitude movements. Experimental Brain Research 189, 49–60. https://doi.org/10.1007/s00221-008-1397-6

Ernst, M.O., Banks, M.S., 2002. Humans integrate visual and haptic information in a statistically optimal fashion. Nature 415, 429–33.

Fengler, A., Govindarajan, L.N., Chen, T., Frank, M.J., 2021. Likelihood approximation networks (LANs) for fast inference of simulation models in cognitive neuroscience. Elife 10.

Fortin, C., 2003. Attentional Time-Sharing in Interval Timing, in: Functional and Neural Mechanisms of Interval Timing. CRC Press. https://doi.org/10.1201/9780203009574.ch9

Hartanto, T.A., Krafft, C.E., Iosif, A.M., Schweitzer, J.B., 2016. A trial-by-trial analysis reveals more intense physical activity is associated with better cognitive control performance in attention-deficit/hyperactivity disorder. Child Neuropsychol 22, 618–26.

Hartcher-O’Brien, J., Luca, M.D., Ernst, M.O., 2014. The Duration of Uncertain Times: Audiovisual Information about Intervals Is Integrated in a Statistically Optimal Fashion. PLoS ONE 9, e89339. https://doi.org/10.1371/journal.pone.0089339

Iwasaki, M., Noguchi, Y., Kakigi, R., 2018. Neural correlates of time distortion in a preaction period. Human Brain Mapping 40, 804–817. https://doi.org/10.1002/hbm.24413

Lemoine, L., Lunven, M., Bapst, B., Cleret, de L.L., de, G.V., Bachoud-Lévi, A.C., 2021. The specific role of the striatum in interval timing: The Huntington’s disease model. Neuroimage Clin 32, 102865.

McAuley, J.H., Marsden, C.D., 2000. Physiological and pathological tremors and rhythmic central motor control. Brain 123 (Pt 8), 1545–67.

Mioni, G., Grassi, M., Tarantino, V., Stablum, F., Grondin, S., Bisiacchi, P.S., 2016. The impact of a concurrent motor task on auditory and visual temporal discrimination tasks. Atten Percept Psychophys 78, 742–8.

Oldfield, R.C., 1971. The assessment and analysis of handedness: The Edinburgh inventory. Neuropsychologia 9, 97–113. https://doi.org/10.1016/0028-3932(71)90067-4

Palmer, J., Huk, A.C., Shadlen, M.N., 2005. The effect of stimulus strength on the speed and accuracy of a perceptual decision. J Vis 5, 376–404.

Pedrosa, D.J., Nelles, C., Maier, F., Eggers, C., Burghaus, L., Fink, G.R., Wittmann, M., Timmermann, L., 2016. Variance of essential tremor patients’ time reproduction deficits. Mov Disord 31, 1428–9.

Petzschner, F.H., Glasauer, S., Stephan, K.E., 2015. A Bayesian perspective on magnitude estimation. Trends in Cognitive Sciences 19, 285–293. https://doi.org/10.1016/j.tics.2015.03.002

Pineda, J.A., 2005. The functional significance of mu rhythms: Translating “seeing” and “hearing” into “doing”. Brain Research Reviews 50, 57–68. https://doi.org/10.1016/j.brainresrev.2005.04.005

Ratcliff, R., 1978. A theory of memory retrieval. Psychological Review 85, 59–108. https://doi.org/10.1037/0033-295x.85.2.59

Ratcliff, R., 1993. Methods for dealing with reaction time outliers. Psychol Bull 114, 510–32.

Rohenkohl, G., Cravo, A.M., Wyart, V., Nobre, A.C., 2012. Temporal expectation improves the quality of sensory information. J Neurosci 32, 8424–8428.

Ross, J.M., Comstock, D.C., Iversen, J.R., Makeig, S., Balasubramaniam, R., 2022. Cortical mu rhythms during action and passive music listening. J Neurophysiol 127, 213–224.

Schütt, H.H., Harmeling, S., Macke, J.H., Wichmann, F.A., 2016. Painfree and accurate Bayesian estimation of psychometric functions for (potentially) overdispersed data. Vision Res 122, 105–123.

Shi, Z., Church, R.M., Meck, W.H., 2013. Bayesian optimization of time perception. Trends Cogn Sci 17, 556–64.

Singh, A., Cole, R.C., Espinoza, A.I., Evans, A., Cao, S., Cavanagh, J.F., Narayanan, N.S., 2021. Timing variability and midfrontal 4Hz rhythms correlate with cognition in Parkinson’s disease. NPJ Parkinsons Dis 7, 14.

Smith, A., Taylor, E., Rogers, J.W., Newman, S., Rubia, K., 2002. Evidence for a pure time perception deficit in children with ADHD. J Child Psychol Psychiatry 43, 529–42.

Vicario, C.M., Martino, D., Spata, F., Defazio, G., Giacchè, R., Martino, V., Rappo, G., Pepi, A.M., Silvestri, P.R., Cardona, F., 2010. Time processing in children with Tourette’s syndrome. Brain Cogn 73, 28–34.

Voss, A., Lerche, V., Mertens, U., Voss, J., 2019. Sequential sampling models with variable boundaries and non-normal noise: A comparison of six models. Psychon Bull Rev 26, 813–832.

Voss, A., Rothermund, K., Voss, J., 2004. Interpreting the parameters of the diffusion model: An empirical validation. Memory &amp Cognition 32, 1206–1220. https://doi.org/10.3758/bf03196893

Wiecki, T.V., Sofer, I., Frank, M.J., 2013. HDDM: Hierarchical Bayesian estimation of the Drift-Diffusion Model in Python. Front Neuroinform 7, 14.

Wiener, M., Thompson, J.C., Coslett, H.B., 2014. Continuous carryover of temporal context dissociates response bias from perceptual influence for duration. PLoS One 9, e100803.

Wiener, M., Zhou, W., Bader, F., Joiner, W.M., 2019. Movement Improves the Quality of Temporal Perception and Decision-Making. eNeuro 6.

van Wassenhove, V., Buonomano, D.V., Shimojo, S., Shams, L., 2008. Distortions of subjective time perception within and across senses. PLoS One 3, e1437.

